# The pseudoenzyme β-amylase9 from Arabidopsis binds to and enhances the activity of α-amylase3: A possible mechanism to promote stress-induced starch degradation

**DOI:** 10.1101/2024.08.07.607052

**Authors:** Christopher E. Berndsen, Amanda R. Storm, Angelina M. Sardelli, Sheikh R. Hossain, Kristen R. Clermont, Luke M. McFather, Mafe A. Connor, Jonathan D. Monroe

## Abstract

Starch accumulation in plant tissues provides an important carbon source at night and for regrowth after periods of dormancy and in times of stress. Both ɑ- and β-amylases (AMYs and BAMs, respectively) catalyze starch hydrolysis, but their functional roles are unclear. Moreover, the presence of catalytically inactive amylases that show starch excess phenotypes when deleted presents an interesting series of questions on how starch degradation is regulated. Plants lacking one of these catalytically inactive β-amylases, BAM9, were shown to have enhanced starch accumulation when combined with mutations in BAM1 and BAM3, the primary starch degrading BAMs in response to stress and at night, respectively. Importantly, BAM9 has been reported to be transcriptionally induced by stress through activation of SnRK1. Using yeast two-hybrid experiments, we identified the plastid-localized AMY3 as a potential interaction partner for BAM9. We found that BAM9 interacted with AMY3 *in vitro* and that BAM9 enhances AMY3 activity 3-fold. Modeling of the AMY3-BAM9 complex revealed a previously undescribed N-terminal structural feature in AMY3 that we call the alpha-alpha hairpin that could serve as a potential interaction site. Additionally, AMY3 lacking the alpha-alpha hairpin is unaffected by BAM9. Structural analysis of AMY3 showed that it can form a homodimer in solution and that BAM9 appears to replace one of the AMY3 monomers to form a heterodimer. Collectively these data suggest that BAM9 is a pseudoamylase that activates AMY3 in response to cellular stress, possibly facilitating starch degradation to provide an additional energy source for stress recovery.

## INTRODUCTION

Starch is a polymer of glucose that accumulates in plant tissues when the energy supply exceeds demand and is broken down later to meet energy needs when the supply of photosynthate is limited (Smith and Zeeman, 2020). In unstressed photosynthetic cells, starch granules are normally synthesized in plastids during the light hours using photosynthate directly and are degraded at night at a rate that is sufficient to nearly exhaust the supply of starch by the following dawn (Gibon *et al*., 2004; Graf *et al*., 2010). In non-photosynthetic cells, starch is synthesized in amyloplasts, typically using sucrose imported from photosynthetic cells, and it acts as an energy buffer to ensure a supply of reduced carbon to fuel future metabolism. Starch degradation also occurs in guard cells at dawn to generate osmotica to aid in stomatal opening (Horrer *et al*., 2016). In addition to these temporally predictable needs, starch metabolism is also altered by a variety of abiotic and biotic stresses to presumably fuel defense strategies (Thalmann and Santelia, 2017; Ribeiro, Stitt and Hotta, 2022). These diverse roles for starch likely necessitate a diverse array of regulatory mechanisms, yet these mechanisms are poorly understood.

Starch granules contain two types of polymers: amylopectin, which comprises ∼80% of the granule, and amylose, which comprises the remainder. Whereas amylose is composed of long chains of ɑ-1,4-linked glucose with occasional short ɑ-1,6-branches, amylopectin is highly branched with layers of short, double-helical chains 12-20 glucose units long that are packed tightly together ultimately creating a very large, osmotically inert granule (Smith and Zeeman, 2020). In leaf mesophyll cells at night, starch degradation commences with phosphorylation of occasional C3 and C6 positions of glucose residues in the outer chains by GLUCAN WATER DIKINASE1 (GWD1) and PHOSPHOGLUCAN WATER DIKINASE (PWD) (Ritte *et al*., 2002; Kotting *et al*., 2005) causing the outer chains to become hydrated and available for hydrolysis (Hejazi *et al*., 2008). Hydrolysis of the ɑ-1,4-bonds at night is accomplished primarily by β-AMYLASE3 (BAM3) and to some extent by BAM1 (Fulton *et al*., 2008; Monroe, 2020). Complete hydrolysis of a chain also requires the removal of phosphates by two phosphatases, STARCH EXCESS4 (SEX4) and LIKE SEX FOUR2 (LSF2) (Kötting *et al*., 2009; Comparot-Moss *et al*., 2010), and debranching by ISOAMYLASE3 (ISA3), and LIMIT DEXTRINASE (LDA) (Niittylä *et al*., 2006; Kötting *et al*., 2009).

The pathway used for starch degradation in leaves during daylight is likely to be somewhat different than nighttime starch degradation (Delatte *et al*., 2006) but it has received less attention. In Arabidopsis guard cells, starch is rapidly degraded at dawn to generate osmotically active metabolites to aid stomatal opening, and it was shown that a plastid-localized ɑ-AMYLASE3 (AMY3) along with BAM1 are involved (Valerio *et al*., 2011; Horrer *et al*., 2016; Flütsch *et al*., 2020). Both AMY3 and BAM1 are also involved in starch degradation in response to osmotic stress (Valerio *et al*., 2011; Thalmann *et al*., 2016). Indeed, AMY3 was identified as a component of chloroplast stress granules (Chodasiewicz *et al*., 2020). These granules form transiently under heat stress and are thought to protect proteins from thermal denaturation and possibly render enzymes inactive quickly (Jain *et al*., 2016; Protter and Parker, 2016). Both BAM1 and AMY3 are activated by reduced thioredoxin, suggesting that they function during the daylight (Sparla *et al*., 2006; Seung *et al*., 2013), and both are transcriptionally induced by abscisic acid supporting their potential role in stress responses (Thalmann *et al*., 2016).

In addition to BAM3 and BAM1, there are seven other BAM genes in Arabidopsis, three of which are not located in plastids (BAM5, BAM7, and BAM8) and are thus unlikely to play a direct role in starch metabolism (Monroe and Storm, 2018). Of the plastid localized forms, BAM1, BAM2, BAM3, and BAM6 are catalytically active, but the functions of BAM2 and BAM6 are unknown. The remaining two, BAM4 and BAM9, lack catalytic activity (Li *et al*., 2009; David *et al*., 2021). In addition to providing evidence that BAM9 is catalytically inactive, David et al. (2021) also noticed from public transcriptome data that a variety of environmental stresses induced the expression of BAM9. Mutants lacking BAM9 had a starch excess phenotype that was enhanced in plants that also lacked BAM1 and BAM3. The authors concluded that BAM9 probably played some role in regulating starch degradation, perhaps under stress (David *et al*., 2021).

The first report on the pattern of BAM9 expression was from Chandler et al. (2001), showing that transcription of *BAM9* (also called *BMY3*) was highest in roots and reproductive structures and that diurnal expression of BAM9 in leaves peaked at the night/day interface, coincidently, the time of lowest energy status (Chandler, Apel and Melzer, 2001; Gibon *et al*., 2004; Graf *et al*., 2010). The spike in BAM9 expression at dawn is also supported by publicly available microarray data (Winter *et al*., 2007). BAM9 was also upregulated in plants overexpressing KIN10, a subunit of SnRK1 (Baena-González *et al*., 2007). SnRK1 is a highly conserved energy sensor in all eukaryotic cells that is activated by low energy status and inhibited by high energy status via trehalose 6-P (Polge and Thomas, 2007; Delatte *et al*., 2011). Recently, Viana et al. (2021) showed that BAM9 is directly induced via its G-Box element by bZIP63 after bZIP63 is activated by the protein kinase SnRK1 (Viana *et al*., 2021). Together, these observations suggest that BAM9 may play a role in regulating the response to low energy status or to stresses requiring heightened energy availability for an effective response.

Given the phenotype of *bam9* mutants and BAM9’s lack of enzymatic function, we suspected that BAM9 acts as a pseudoenzyme, regulating starch degradation through an unknown mechanism. We screened for interaction partners to explore this potential function for BAM9 and identified the ɑ-amylase, AMY3, as a possible interaction partner. We found that BAM9 forms a 1:1 complex with AMY3, which enhances the starch-degrading activity of AMY3. The interaction between BAM9 and AMY3 is mediated through the N-terminus of AMY3, likely through a previously undescribed region consisting of an alpha-alpha hairpin. Surprisingly, AMY3 alone formed a homodimer that had lower activity than AMY3 in the presence of BAM9. Collectively, the data support the role of BAM9 as a pseudoamylase in derepressing AMY3 activity.

## METHODS

### Yeast-Two-Hybrid screening

#### Generation of pAS1-BAM9 transformed Y190 cells

Yeast Two-Hybrid experiments were performed using the Walker Two-hybrid cell stocks and cDNA library (Arabidopsis Biological Resource Center [ABRC] stock no. CD4-10). Reporter strain Y190 yeast cells, having *LacZ* and *His3* reporter genes, were transformed with a pAS1-BAM9 bait plasmid containing the BAM9 cDNA cloned in frame with the GAL4 DNA-binding domain. A modified version of the protocol by Tian et al. (2012) was used for a small-scale transformation, with 100 μL Y190 cells resuspended in Solution I (100 mM LiAc, 10 mM Tris-HCl [pH 7.5], 1 mM EDTA) combined with 2 μL of pAS1-BAM9 DNA (0.25 μg/μL), 10 μL of boiled calf thymus DNA (10 mg/mL) and 700 μL Solution II (100 mM LiAc, 10 mM Tris-HCl [pH 7.5], 1 mM EDTA, and 50% polyethylene glycol [PEG]-3350) (Tian, Zhao and Liu, 2012). Transformed cells were resuspended in 200 μl sterile deionized water and serial dilutions (1:4 and 1:100) were plated on Complete Media lacking tryptophan (CM -Trp; Difco Yeast Nitrogen Base without amino acids, 2% glucose, 4 mM serine, 0.1 mM adenine hemisulfate salt, 0.15 mM lysine HCl, 0.3 mM phenylalanine, 0.15 mM tyrosine, 0.1 mM arginine HCl, 0.15 mM methionine, 1.5 mM threonine, 0.75 mM leucine, and 0.1 mM histidine). The resulting pAS1-BAM9 strain was tested for self-activation by growth on CM -Trp -His plates with increasing concentrations (0 – 75 mM) of 3-amino-1,2,4-triazole (3-AT), a competitive inhibitor of the *HIS3* reporter gene product, and were found to be unable to grow on CM -Trp -His plates when supplemented with 25 mM or more of 3-AT. All subsequent selection plates were made with 25 mM 3-AT.

#### cDNA library screen

The pAS1-BAM9 strain was used to screen a library of prey vectors. The *A. thaliana* Walker cDNA library, made with random-primed mRNA isolated from mature Arabidopsis leaves and roots, was amplified and excised into a pACT plasmid in fusion with the sequence encoding the GAL4 activation domain according to the ABRC manual provided with the library. The cDNA library screen was performed as described in Tian et al. (2012). Y190 pAS1-BAM9 yeast cells were grown in 500 mL YPD media to a density of 0.4 OD_600_, resuspended in 2.5 mL Solution I and combined with 30 µg purified cDNA library, 125 μL of boiled calf thymus DNA, (10 mg/mL) and 15 mL of Solution II. After transformation, cells were spread equally on a total of 46 selection plates with Complete Media lacking tryptophan, leucine, and histidine supplemented with 25 mM 3-AT (CM -Trp -Leu -His +3-AT plates), and growth was monitored over 1-3 weeks. Colonies that were able to grow in the absence of histidine, indicating expression of the *His3* reporter gene, were also tested with an X-gal assay as described in Tian et al. (2012). Colonies that grew on the -His plates were grown on CM-Trp -Leu +3-AT plates and colonies were transferred to a nitrocellulose membrane, lysed through freezing with liquid nitrogen, incubated with 20 µL of X-gal (20 mg/mL in DMF) in 1.5 mL of Buffer Z (60 mM Na_2_HPO_4_, 40 mM NaH_2_PO_4_, 10 mM KCl, 1 mM MgSO_4_, pH 7.0), and screened for the formation of blue colonies over 24 hours. This cDNA library screen was performed twice.

#### Isolation of prey vectors

Colonies showing evidence of both *LacZ* and *His3* expression were cultured and lysed using Zymolyase solution (50:49:1, glycerol:Zymolyase [G-Biosciences]: 1M Tris) and prey plasmids were purified by alkaline lysis miniprep (Joly, 1996). Plasmids were transformed into *E. coli* cells (DH5α) using carbenicillin selection and streaked for isolation to create transformed prey stocks. Mini-prepped prey plasmids were sequenced using the Gal4AD primer (5’-TACCACTACAATGGATG-3’). The prey sequences were identified using NCBI BLASTn and the UniProt database was used to identify aspects of the identified protein such as location, family and function.

Isolates representing the top protein hit, AMY3, were further evaluated to test for false positives. Purified prey plasmids were transformed into 1) Y190 yeast with no plasmids, 2) Y190 yeast with empty pAS1, 3) Y190 yeast with pAS1-BAM9, and 4) Y190 yeast with a negative control bait vector pAS1-SNF1. These transformants were grown on CM –Leu –His +3-AT media in the case of the Y190 transformants or CM –Leu –Trp –His +3-AT media in the case of transformants containing pAS1. Growth on -HIS media was observed, and a X-gal assay was conducted on each plate. Only the pAS1-BAM9 transformants produced viable X-Gal positive colonies.

### Models of BAM9 and AMY3 and complexes

Models of BAM9, AMY3, a 1:1 complex of BAM9 and AMY3, and the AAH domain of AMY3 were generated using AlphaFold3 (Abramson *et al*., 2024). Sequences with the chloroplast targeting sequence were downloaded from UniProt (BAM9:Q8VYW2; AMY3: Q94A41) and provided as inputs for AlphaFold3. Sequences were submitted in a 1:1 ratio several times with a different seed number. No information on the putative interaction site was provided.

### Protein expression in *E. coli* and purification

Initially, we purified AMY3 from *E. coli* using the construct from Seung, et al. (Seung *et al*., 2013). Following issues with AMY3 degradation, we synthesized AMY3 lacking the chloroplast transit peptide (CTP) in pET21a (Genscript) and expressed the construct in BL21-CodonPlus *E. coli* cells. We then purified AMY3 in a single day and found significant degradation of the protein began within 12-18 hours. Cells expressing AMY3 were grown in 0.06 mg/mL ampicillin to an OD_600_ of approximately 0.6 at 37 °C and 225 rpm, then induced with 1 mM Isopropyl-β-D-thiogalactoside (IPTG) at 20 °C overnight. Cells were lysed in 50 mM Tris, pH 8.0, 100 mM sodium chloride, 0.1 mM EDTA, 1 mM Imidazole, and an EDTA-free protease-inhibitor tablet (Pierce A32965) using sonication at 65% power for 5 seconds on and 5 seconds off for 5 minutes. Cell debris was removed by centrifugation at 17,000 rcf at 4 °C. Supernatant was then poured over 2 mL of Ni-NTA resin, and the column was allowed to drain via gravity. The column was washed with 20 mL of Wash Buffer (50 mM Tris, pH 8.0, 300 mM sodium chloride, 0.1 mM EDTA, 10 mM Imidazole). Protein was eluted with 10 mL steps from 9:1 wash buffer:elution buffer (50 mM Tris, pH 8.0, 50 mM sodium chloride, 0.1 mM EDTA, 300 mM Imidazole) to 0:10 wash buffer:elution buffer. The presence of protein was confirmed via SDS-PAGE, combined elutions were concentrated to 2 mL, and then injected onto a Sephacryl-200 16/60 column in 50 mM sodium phosphate, pH 7, 100 mM sodium chloride, and 5 mM DTT. The presence of protein in the fractions was confirmed via SDS-PAGE, protein-containing fractions were pooled, concentrated, and then stored at −80 °C. The AMY3 amylase catalytic domain (amino acids 484-837) was cloned after codon optimization into pET21a, expressed, purified, and stored as described for full-length AMY3 above.

BAM9 was expressed in BL21 cells with pETDuet-1-BAM9-B cultured with 0.06 mg/mL ampicillin. Cells were grown to an OD_600_ of approximately 0.6 at 37 °C and 225 rpm, then induced with 1 mM Isopropyl-β-D-thiogalactoside (IPTG) at 20 °C overnight. Cells were pelleted, frozen, and then sonicated on ice in 40 mL Buffer A (0.5 M NaCl, 50 mM NaH_2_PO_4_, 0.2 mM TCEP, and 2 mM imidazole, pH 8) with an EDTA-free protease-inhibitor tablet (Pierce A32965). Cell debris was removed by centrifugation at 17,000 rcf at 4 °C. This supernatant was then diluted to 175 mL with Buffer A. Protein was then purified using an AKTA Start protein purification system with a 1 mL TALON cobalt-affinity column using washes of 2%, 10%, 20%, and 100% Buffer B (Cytiva Life Science, Marlborough, Massachusetts, USA). Buffer B contained 0.5 M NaCl, 50 mM NaH_2_PO_4_, 0.2 mM TCEP, and 200 mM imidazole, pH 8. Buffers A and B were supplemented with 3 mM benzamidine (TCI) when purifying AMY3 to reduce degradation. The BAM9 that was analyzed using small-angle X-ray scattering (SAXS) was additionally purified using size exclusion chromatography (SEC) with a Superdex 100 column (Cytiva) in Buffer A.

### Amylase enzyme assays

Amylase activity of purified AMY3 and BAM9 were assayed in 0.5 mL assay solution containing 50 mM HEPES (pH 8) with 1 mM MgCl, 0.2 mM TCEP, and 20 mg/mL soluble starch (Acros Organics), amylopectin, or amylose as the substrate. Soluble starch and amylopectin were dissolved by heating, whereas amylose was dissolved in 2 N KOH, and then neutralized with 2 N HCl. Proteins were diluted in 50 mM MOPS, pH 7, with 1 mg/mL bovine serum albumin. Assays were initiated with the addition of enzyme, incubated for 20 minutes at 25 °C, and then boiled for three minutes to denature the proteins. Reducing sugars were quantified using the Somogyi-Nelson assay (Nelson, 1944) with maltose as the standard. For some assays, indicated concentrations of purified BAM9 were added to the diluted AMY3 prior to the initiation of the assays.

### Protein-protein crosslinking

BAM9 and AMY3 were mixed to a final concentration of 10 µM for both proteins in 20 mM HEPES, pH 7.5. Proteins were pre-incubated at room temperature (∼22 ℃) in 25 µL for 15 minutes before adding 30 µL of 0.05% glutaraldehyde. Crosslinking occurred for 30 minutes at room temperature before being quenched with 30 µL of 1 M Tris base. Glutaraldehyde and Tris were allowed to react for 5 minutes before the reaction was mixed with 2x SDS-PAGE loading dye. Proteins were then separated via SDS-PAGE.

### Small-angle X-ray scattering of BAM9 and AMY3

Purified protein samples of AMY3 and BAM9 alone and combined were shipped overnight at 4 °C and analyzed using SEC-SAXS at the SIBYLS beamline 12.3.1, Advanced Light Source, Lawrence Berkeley National Laboratory, Berkeley, CA (Classen *et al*., 2013; Rosenberg, Hura and Hammel, 2022). Samples were separated on either a Shodex KW-803 column at a flow rate of 0.5 mL/min at 10 °C, and eluate was measured in-line with UV/vis absorbance at 280 nm, Multi-Angle X-ray Scattering (MALS), and SAXS. The incident light wavelength was 1.127 Å at a sample-to-detector distance of 2.1 m. This setup results in scattering vectors, q, ranging from 0.0114 Å^-1^ to 0.4 Å^-1^, where the scattering vector is defined as 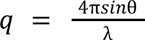 and 2θ is the measured scattering angle. Raw data are available from Simple Scattering under codes XS5XWKRW for BAM-AMY3, XS1RR5P9 for BAM9, and XSH2NT1L for AMY3 (Kim *et al*., 2024).

Radially averaged SAXS data files were processed and analyzed in Scatter IV and RAW (Hopkins, 2024). Peaks containing potential AMY3-BAM9 complexes were identified by comparing plots of integrated intensity vs. elution volume to similar plots for BAM9 and AMY3 alone. For BilboMD fitting of structural ensembles, the top structure and PAE from AlphaFold3 were provided to generate the file of flexible regions, followed by modeling through the standard BilboMD workflow (Pelikan, Hura and Hammel, 2009; Chinnam *et al*., 2023). Processed SAXS data and models are available on the SASBDB with codes SASDVW5 (AMY3), SASDVX5 (BAM9), and SASDVY5 (BAM9+AMY3 (Kikhney *et al*., 2019).

### Protein sequence analysis

Amino acid sequences of Arabidopsis BAM9 (NP_197368.1) and AMY3 (NP_564977.1) were used to conduct BLASTp searches at NCBI RefSeq database (blast.ncbi.nlm.nih.gov/), Ginkgo database (ginkgo.zju.edu.cn/genome/), MarpolBase (marchantia.info/), and CNGBdb transcriptome database (db.cngb.org). Sequences were aligned using Clustal Omega (Madeira *et al*., 2019) and analyzed manually before mapping onto a phylogenetic tree of selected Viridiplantae generated using TimeTree (Kumar *et al*., 2017).

## RESULTS

### Yeast Two-Hybrid screen for BAM9 interactors

BAM9 is a catalytically inactive member of the β-amylase family with a proposed regulatory role in starch metabolism (David *et al*., 2021). To identify binding partners of BAM9 that may suggest potential cellular functions, an Arabidopsis yeast two-hybrid cDNA library was screened using BAM9 as bait. Around 10^6^ transformants were generated, which resulted in 105 colonies able to grow on -HIS media, 74 of which also showed evidence of *LacZ* activity. Sequencing of these isolates identified 21 unique gene hits, 8 of which are predicted or known to be localized to the chloroplast along with BAM9 (Supplementary Table S1). Of these potential chloroplast interactors, 3 unique, independent prey plasmids contained cDNA encoding a portion of the catalytically active α-amylase AMY3. The Arabidopsis genome contains three AMY genes but only AMY3 encodes a plastid-targeted protein (Yu et al., 2005). Unlike AMY1 and AMY2, which encode only a catalytic domain, AMY3 is about twice as long as the others and encodes two carbohydrate-binding modules (CBM) in the CBM45 family at the N-terminus and a long spacer between the second CBM and catalytic domain (Seung *et al*., 2013). Within this spacer, we predicted a previously unannotated alpha-alpha-hairpin (AAH) structure. Intriguingly, all the AMY3 prey plasmids had sequences containing the AAH structure (Fig. 1).

**Figure 1.**
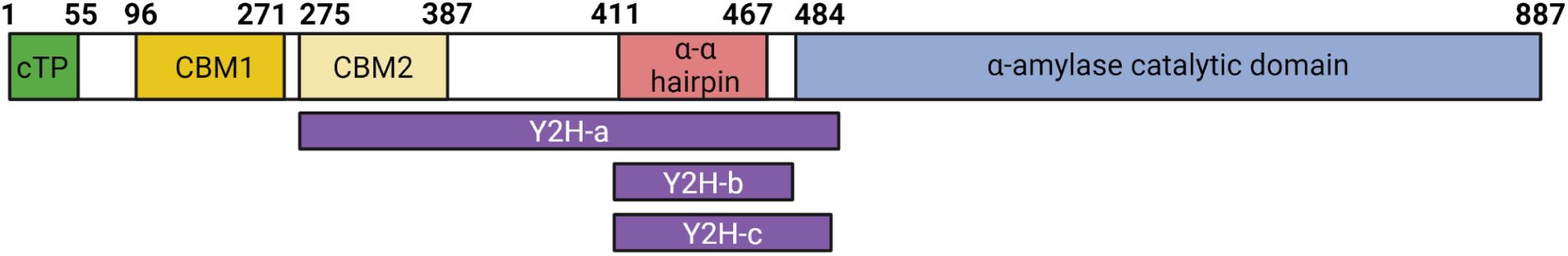
Domain organization of Arabidopsis AMY3. The numbers indicate the amino acid boundaries for each region as defined by Uniprot (Q94A41) and the AlphaFold3 model. cTP stands for chloroplast transit peptide, which is removed after AMY3 is transported to the chloroplast. CBM, Carbohydrate binding modules CBM45. Purple Y2H bars indicate the prey plasmid sequences identified as interacting with BAM9 in the yeast two-hybrid experiment. A complete list of possible BAM9 plastidic interaction partners are found in Supplementary Table S1.

### BAM9 enhances AMY3 activity

To confirm the interaction of AMY3 and BAM9, we purified recombinantly expressed versions of both proteins. BAM9 was purified as a homogenous protein, while AMY3 showed some signs of degradation even in the presence of protease inhibitors and when purified to apparent homogeneity (Fig. 2A). In addition, we observed that AMY3 migrated on SDS-PAGE at a higher molecular weight (∼110 kDa) than predicted based on sequence (95 kDa). Proteins with disordered regions are known to display this behavior along with degradation (Habchi *et al*., 2014).

**Figure 2.**
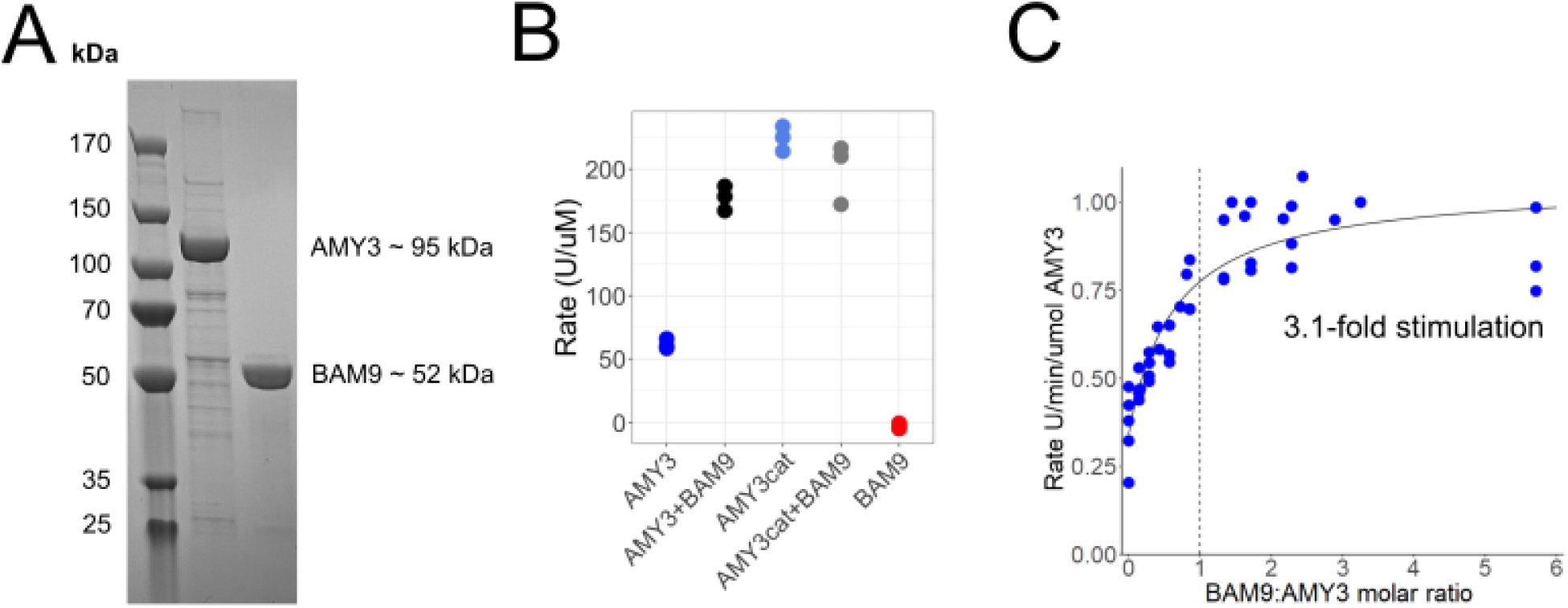
Purification and amylase activity of BAM9 and AMY3. **(A)** SDS-PAGE of recombinant AMY3 and BAM9 after nickel affinity and size-exclusion chromatography purification. Sequence-based molecular weights for each protein are indicated. AMY3 consistently migrates slower than expected based on the sequence weight. This phenomenon has been observed by others studying AMY3 (Seung *et al*., 2013). **(B)** Stimulation of AMY3 activity by BAM9. AMY3_cat_ is a purified, recombinant truncation of AMY3, which includes only the ɑ-amylase domain. Data shown are representative data from three assays and the raw scatter of the measurements from this experiment is shown. **(C)** Titration of BAM9 to estimate the stoichiometry of the BAM9-AMY3 complex. Dashed vertical line indicates a 1:1 molar ratio, and combined data from three experiments are shown in the plot.

We then performed starch hydrolysis assays with each protein alone and in a 1:1 (molar) mix of both proteins. When assayed separately, AMY3 was active with soluble starch as the substrate, but we detected no activity when BAM9 alone was used (Fig. 2B), consistent with BAM9 being a catalytically inactive pseudoenzyme (David *et al*., 2021). To determine whether the presence of BAM9 affected the activity of AMY3, we combined the two proteins prior to initiating assays. The activity of AMY3 was about 3-fold higher in the presence of equal molar concentrations of BAM9 than with AMY3 alone (Fig. 2B). A similar increase was also seen with amylopectin and amylose substrates (Supplementary Fig. S1A). We also tried changing the salt in the buffer to potassium given our previous findings that K^+^ stimulates the activity of BAM2, finding that K^+^ had no effect on AMY3 activity or the activation by BAM9 (Supplementary Fig. S1B) (Sholes *et al*., 2024).

Next, we titrated BAM9 into a fixed concentration of full-length AMY3 to determine whether AMY3 was saturable. This would suggest that AMY3 and BAM9 form a discrete complex and indicate the stoichiometry of BAM9 to AMY3. We observed that the plateau in activity begins at ∼ a 1:1 ratio of BAM9 to AMY3 and that the max stimulation of activity is about threefold (Fig. 2C).

Interestingly, after size exclusion chromatography, when AMY3 was purified without the addition of benzamidine, we noticed that AMY3 became degraded to a truncated 45 kDa protein (tAMY3), consistent with the size of the catalytic domain. This tAMY3 was active, as previously observed (Yu *et al*., 2005), but the activity was not stimulated by BAM9, suggesting that the carbohydrate-binding domains (CBMs) and/or alpha-alpha hairpin (AAH) are needed for BAM9 to interact with and activate AMY3 (data not shown). Following this result, we optimized the purification of AMY3 to reduce degradation and created a clone containing only the AMY3 amylase domain (residues 484-837), which we refer to as AMY_cat_. When we tested the stimulation of this construct by BAM9, we found that BAM9 did not stimulate AMY3_cat_, as we observed with the degradation product, tAMY3 (Fig. 2B). Further, the AMY3_cat_ construct alone appears to be more active than full-length AMY3 even in the presence of BAM9. These data suggest that the N-terminus of AMY3 is critical for both regulation of AMY3 and stimulation by BAM9. Collectively, these data suggest that BAM9 forms a complex with AMY3 to stimulate the amylase activity of the complex and that this interaction requires the N-terminal portions of AMY3.

### Dimer model of AMY3-BAM9

Given that there are no experimentally derived structures of BAM9 or AMY3, it was challenging to describe the mechanism of BAM9 stimulation of AMY3 activity. However, we took advantage of the recently released AlphaFold3 to model a heterodimer of BAM9-AMY3 (Abramson *et al*., 2024). The only inputs we provided to the program were the primary sequence of both proteins. The model of BAM9 was consistent with known BAM structures with a pTM score of 0.88, a score indicating high confidence in the model (Adachi *et al*., 1998; Vallée *et al*., 1998; Xu and Zhang, 2010; Sun, Palayam and Shabek, 2022). For the AMY3-BAM9 complex, we produced 10 models to compare and develop a consensus. The highest scoring result had an ipTM score of 0.74, which is in the “grey zone” (ipTM score between 0.6 and 0.8), where the model should be confirmed with additional evidence (Figs. 3A and 3B) (Abramson *et al*., 2024). The predicted alpha-alpha hairpin in AMY3 was also consistently shown to be helical, and 9 out of 10 were modeled as two helices in a hairpin. The second helix in the structure was predicted more confidently than the first helix. Ten models of BAM9 bound to full-length AMY3 were then aligned using the position of BAM9 in order to visualize the various positions of the AMY3 domains relative to BAM9. Whereas the amylase and CBM domains of AMY3 adopted several positions on the surface BAM9, the alpha-alpha hairpin was consistently bound to the same surface region of BAM9 (Fig. 3B). Focusing on the interaction with the alpha-alpha hairpin consistently showed that the second helix in the hairpin (dark blue in Fig. 3C) was modeled bound to BAM9 at an interaction surface that is conserved among BAM9 sequences from 21 orders of Angiosperms (species and accession numbers are listed in Supplementary Table S2) (Fig. 3C). The sequence of the second helix in the alpha-alpha hairpin of AMY3 appears to be more highly conserved than the first helix among AMY3 sequences from the same 21 orders of Angiosperms (Fig. 3C). The position accuracy error plot generated by AlphaFold 3, which graphically describes the confidence of the position of each amino acid in the model to every other amino acid, further supports this interaction by showing a distinct band of amino acids with lower position error in the predicted interaction site (Supplementary Figure S2). These models suggest that the alpha-alpha hairpin within the N-terminus of AMY3 is a potential interaction site with BAM9.

**Figure 3.**
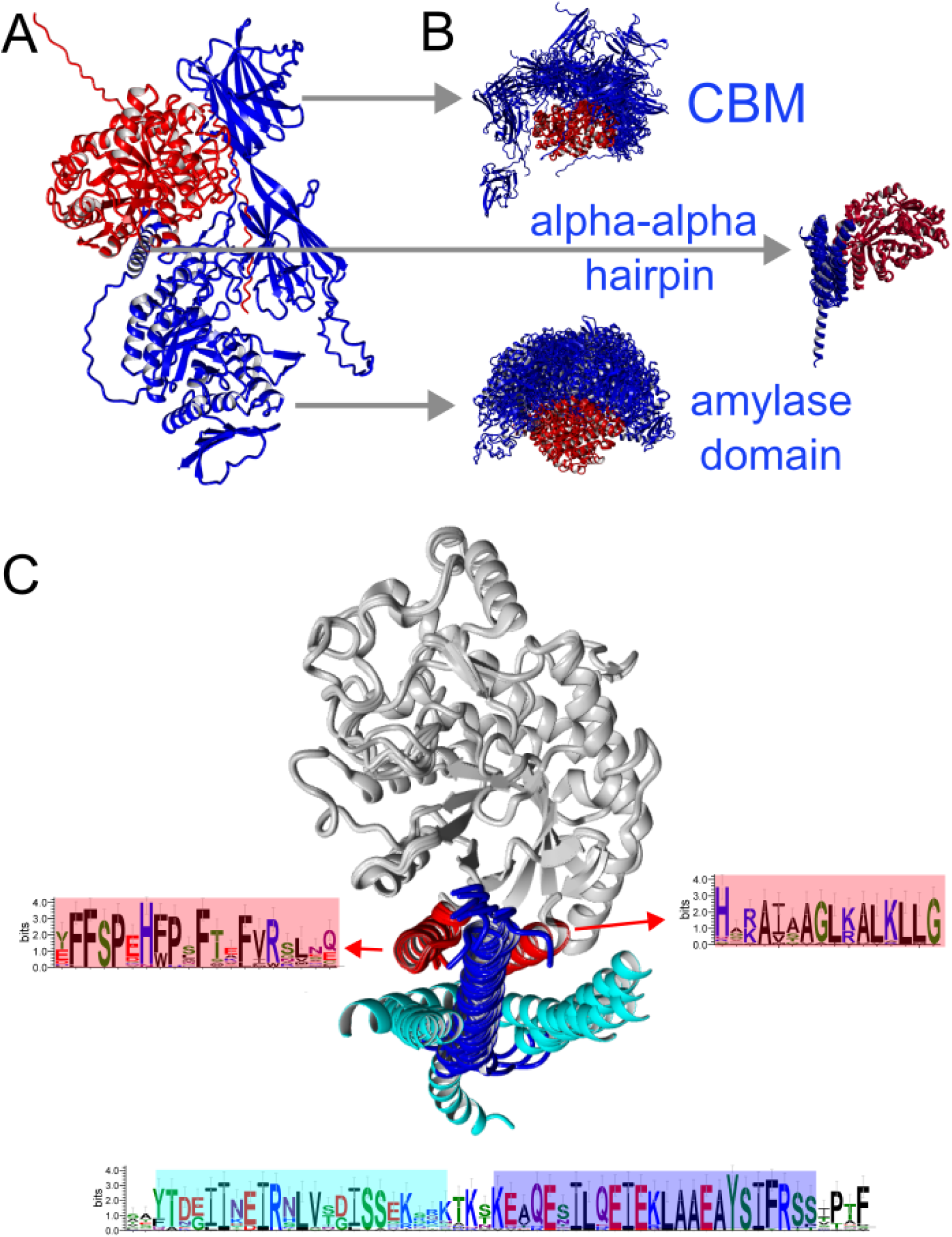
Structural Model of BAM9-AMY3 and conservation of the potential interaction site. **(A)** Top scoring AlphaFold3 model of AMY3 (blue) and BAM9 (red). **(B)** Positioning of individual domains of AMY3 on BAM9 (right) from the 10 AlphaFold models. The top panel shows the positions of the tandem CBM domains, middle is the alpha-alpha hairpin domain, and bottom is the amylase domain (all in blue) relative to BAM9 (red) in alignments of potential AMY3-BAM9 complexes. **(C)** Focus on the 10 positions of the alpha-alpha hairpin (cyan/blue) in the predicted interaction with BAM9 (red/white). The WebLogo of the alpha-alpha hairpin is indicated with cyan or blue rectangles to indicate the first and second respective helix in the hairpin. The two helices within BAM9 and the corresponding WebLogos showing sequence conservation are highlighted in red. WebLogos for AMY3 and BAM9 sequences were generated from alignments of sequences from 21 orders of Angiosperms (see Supplementary Table S2 for species and accession numbers).

### Crosslinking of BAM9 to AMY3

Having found evidence for a functional connection between AMY3 and BAM9, we next attempted to purify an AMY3-BAM9 complex to propose a mechanism of BAM9 activation of AMY3. Initial attempts to co-purify AMY3 and BAM9 as a complex by SEC column chromatography resulted in a broad peak suggesting a diverse set of species in solution. We then turned to glutaraldehyde crosslinking to trap BAM9 and AMY3 in a complex. Unexpectedly, AMY3 formed a multimer(s) when crosslinked without BAM9 (Fig. 4A). The AMY3 monomer appears to run between the 100 and 130 kDa markers on the gel, while the crosslinked species migrates at ∼170 kDa, suggestive of a dimer of AMY3. In contrast, BAM9 alone showed limited multimer formation. When the two proteins were combined and crosslinked, the AMY3 homomultimer was disrupted and replaced by a faster migrating complex with a molecular weight between 130 kDa and 170 kDa, consistent with a heterodimer of BAM9 and AMY3 (147 kDa based on amino acid sequences) (Fig. 4A).

**Figure 4.**
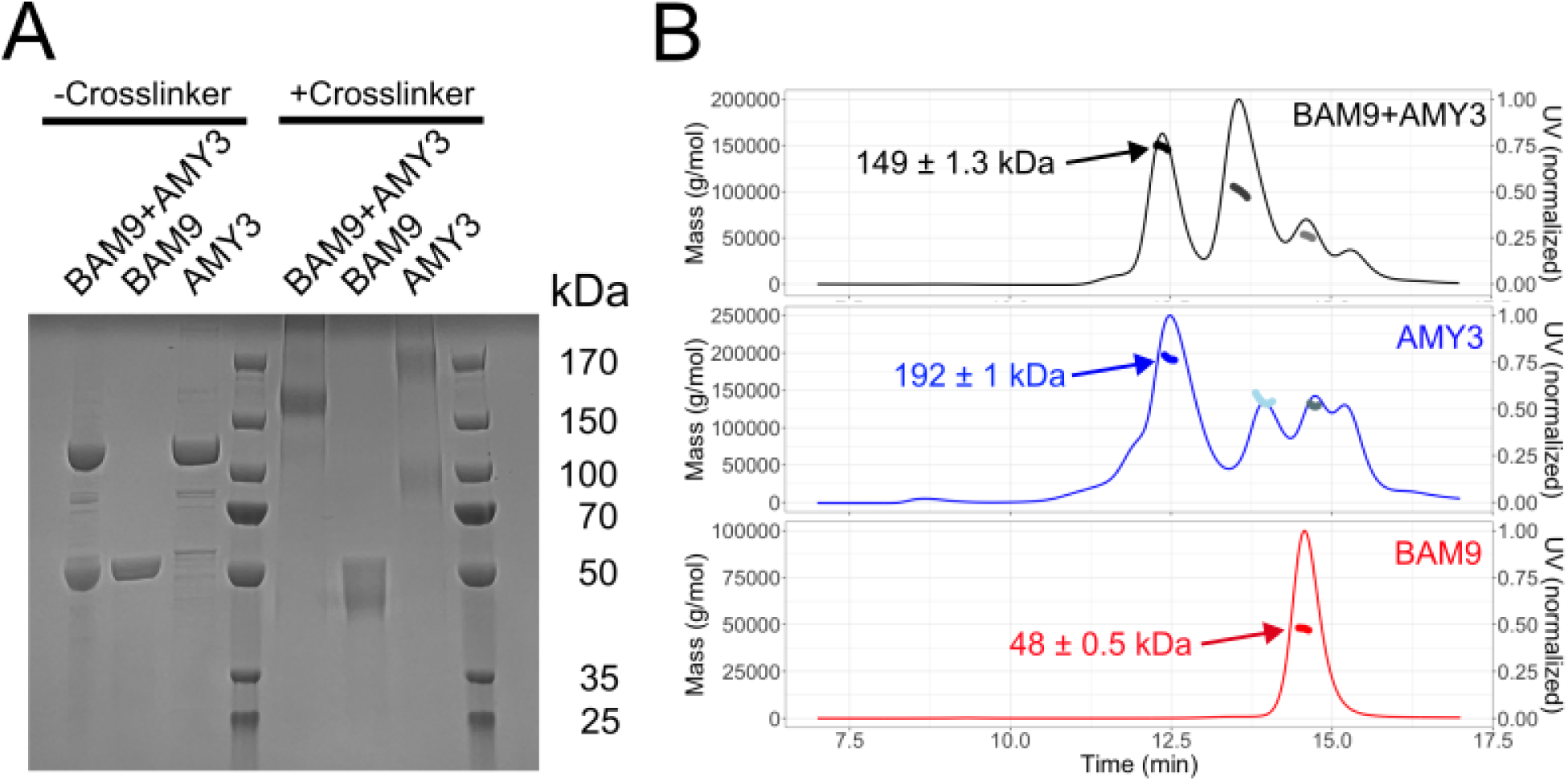
Isolation of the BAM9-AMY3 complex and AMY3 homodimer. **(A)** SDS-PAGE of BAM9, AMY3, and BAM9 with AMY3 alone or after crosslinking with glutaraldehyde. **(B)** Size-exclusion chromatography coupled to multi-angle light scattering (SEC-MALS) of BAM9 (red), AMY3 (blue), and BAM9 with AMY3 (black). Molecular weight measurements from MALS are indicated in points and correspond to the masses indicated on the left-side y-axis.

The diversity of crosslinked complexes explains the broad peak observed in SEC experiments to purify the complex and suggests a dynamic interaction between the AMY3 monomers, dimers, and BAM9 under these conditions (Fig. 4B). To support the crosslinking data and characterize the structure of the BAM9-AMY3 complex, we collected SAXS and MALS data on AMY3, BAM9, and crosslinked BAM9-AMY3 after separation by size exclusion chromatography. The data on BAM9 were consistent with a homogenous, monomeric protein with a molecular weight from MALS of 48 ± 0.5 kDa (Fig. 4B). The data on AMY3 showed three prominent species with MALS molecular weights of 192 ± 1 kDa, 135 ± 1 kDa, and 130 ± 1.3 kDa. Because multidomain proteins can adopt many conformations in solution, the width of the first peak is wide and reflects a diversity of potential radii (Habchi *et al*., 2014). We propose that the latter peaks are complexes of full-length AMY3 with AMY3 degradation products.

Having characterized the solution behavior of AMY3 and BAM9 alone, we next collected data on BAM9 and AMY3. Like the AMY3 sample, the BAM9-AMY3 sample was apparently heterogeneous, and the elution profile of the complex was nearly identical to that of AMY3 (Fig 4B). Alignment of the elution profiles, however, showed that in the BAM9-AMY3 sample, there was an increase in the earliest elution peak time (suggesting a smaller molecular weight complex) mirroring the crosslinking gel where increased BAM9 concentration led to the appearance of a faster migrating band. Since these samples were not crosslinked, this supports an authentic interaction between BAM9 and AMY3 and not a spurious side reaction. The molecular weight from MALS for peak 1 was 149 ± 1.3 kDa, corresponding to a dimer of BAM9-AMY3 with a theoretical molecular weight of 147 kDa (Fig 4B). These data are consistent with the SDS-PAGE gel of crosslinked protein in Figure 4A.

### Evidence for complex formation by SAXS

From the SEC-MALS data, we then analyzed the SAXS data from the peaks with the appropriate molecular weight to develop structural models of the proteins and complexes (Supplementary Table S3). As expected, BAM9 was a compact globular protein according to Kratky plotting of the SAXS data (Fig 5A). Further, we fitted the scattering data to an AlphaFold3 model of BAM9, and the fit had an *X*^2^ of 1.02 (Fig 5B). The first peak in the AMY3 elution during SAXS corresponds to a dimer of full-length AMY3, and Kratky analyses suggested the protein was more flexible relative to BAM9 as indicated by the more gradual decline of the data at higher qRg values (Fig 5A). We then analyzed the first peak to describe the structure of the AMY3 dimer (Supplementary Table S3). Because there is no published structure of AMY3 and it was potentially a flexible protein, we modeled the structure of an AMY3 dimer using AlphaFold3 and then used BilboMD to model potential structural ensembles (Pelikan, Hura and Hammel, 2009; Abramson *et al*., 2024). All of the domains in AMY3 were treated as rigid bodies, and the unstructured linkers were allowed to expand and contract to produce a pool of structures that were fitted to the data to produce a best-fit ensemble. The resulting minimal ensemble consisted of at least two components (X^2^ = 0.8), one compact (∼40%) and one extended (∼60%) (Fig 5C). Adding additional structures to the ensemble did not improve the fit. These data support the dynamic behavior of AMY3 suggested by crosslinking and SEC-MALS experiments. Analysis of the Kratky plots of the AMY3 SAXS data from the first peak and the first peak of the BAM9-AMY3 SAXS data shows differences between the data sets suggestive of a distinct protein complex shape (Fig. 5A). Given the ambiguity of SAXS and the low resolution of the current data, we did not model the BAM9-AMY3 complex further. Overall, the SAXS data are consistent with the formation of a BAM9-AMY3 heterodimer after combining an AMY3 homodimer with BAM9.

**Figure 5.**
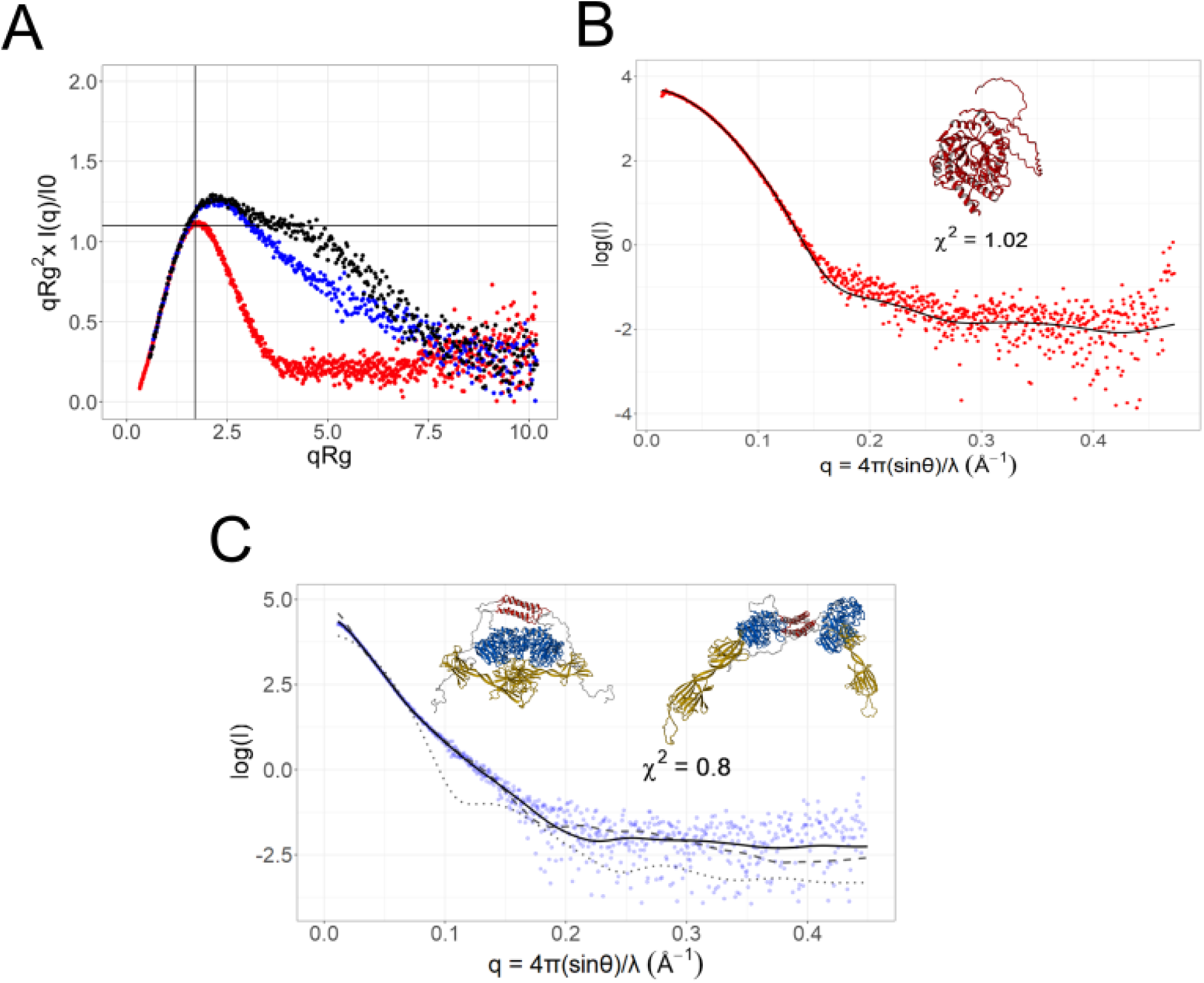
Small-angle X-ray scattering of BAM9, AMY3, and BAM9-AMY3. **(A)** Kratky plots of BAM9 (red), AMY3 (blue), and BAM9 with AMY3 (black) SAXS data. The crossed lines show the “ideal” maximum for globular proteins. Deviations up and to the right of this intersection indicate proteins that are flexible in solution. **(B)** log(intensity) vs. q plot showing the data and the fit of an AlphaFold3 model of BAM9 to the data. **(C)** log(intensity) vs. q plot showing the data and the fit of an AlphaFold3 model of AMY3 to the data. The solid line is the fit of the BilboMD fit to the SAXS data. Dotted and dashed lines indicate the fit of the AlphaFold3 model and a BilboMD-optimized single structure fit to the data, which has X^2^ values of 16 and 2, respectively. The two structures are the BilboMD optimized ensemble that fitted best to the data.

### Evolution of BAM9 and AMY3

Although BAM9 and AMY3 appear to function as a heterodimer in Arabidopsis and probably other flowering plants, each protein may have had an independent function, which converged on the present function within the heterodimer. If they had independent functions, this would be indicated by identifying taxa in which only one of the two genes is present and/or if one of the two genes originated earlier than the other. Monroe et al. (2017) identified Angiosperms as the earliest occurrence of BAM9, but this was based only on the NCBI RefSeq collection of complete genome sequences, which limited the diversity of species analyzed (blast.ncbi.nlm.nih.gov) (Monroe *et al*., 2017). Here, we extended this earlier analysis by searching the NCBI RefSeq database, Ginkgo Database (https://ginkgo.zju.edu.cn/genome/), MarpolBase (https://marchantia.info/), and also the extensive CNGBdb transcriptome database (db.cngb.org). Sequences were identified as BAM9 and AMY3 based on having amino acid sequences and domain structures more similar to Arabidopsis BAM9 and AMY3 than to any other Arabidopsis BAM or AMY. We defined AMY3 sequences as containing the N-terminal CBM and AAH regions, and BAM9 sequences were defined by the lack of the catalytically important flexible loop: “GGNVGD” (David et al., 2021) and containing the highly conserved C-terminal “FFSPEH” sequence shown in Figure 3C that is lacking in closely related BAM1 and BAM3. Sequence identification numbers are listed in Supplementary Table S4.

The presence or absence of putative *BAM9* and *AMY3* genes were mapped onto a phylogenetic tree of Viridiplantae generated by TimeTree5 (timetree.org). Despite possessing several *BAM* and *AMY* genes each, none of the green algae or nonvascular plants that we examined contained a gene resembling *BAM9* or *AMY3* (Fig. 6). Glaring et al. (2011) noted the presence of genes resembling AMY3 in several green algae including *Micromonas pusilla* and *Ostreococcus lucimarinus (Glaring et al., 2011)*. However, these proteins lack the conserved AAH domain. Thus, the earliest occurrence of BAM9 and AMY3 containing the AAH domain is in the pteridophyte *Equisetum hyemale*, suggesting that both genes evolved approximately 400 mya and may have functioned as a heterodimer from their origins. However, in the other three groups of pteridophytes, either one or both genes were apparently lost. All of the ferns in the eusporangiate and leptosporangiate orders that we examined contained *BAM9* but did not contain *AMY3* (Fig. 6). In these taxa, the absence of AMY3 suggests that BAM9 may have an additional function beyond the one described in this study. In the whisk fern, *Psilotum nudum,* and grape fern, *Ophioglossum vulgatum*, both *BAM9* and *AMY3* genes were lost. Nearly all the seed plants examined contained both *BAM9* and *AMY3* with the exception of the Gnetophyta division of gymnosperms, which also lacks both *AMY3* and *BAM9* (Fig. 6). Using AlphaFold 3 we then modeled heterodimers from one member of each of the major groups of land plants possessing both genes, and observed that AMY3 may form a heterodimer with BAM9 similar to what we observe here with the Arabidopsis homologs (Supplementary Figure S3). Thus, we find that BAM9 regulates the starch hydrolysis activity of AMY3 through complex formation and this complex is probably found in many vascular plant species. These data suggest that pseudoamylase regulation of starch hydrolysis is a conserved process.

**Figure 6.**
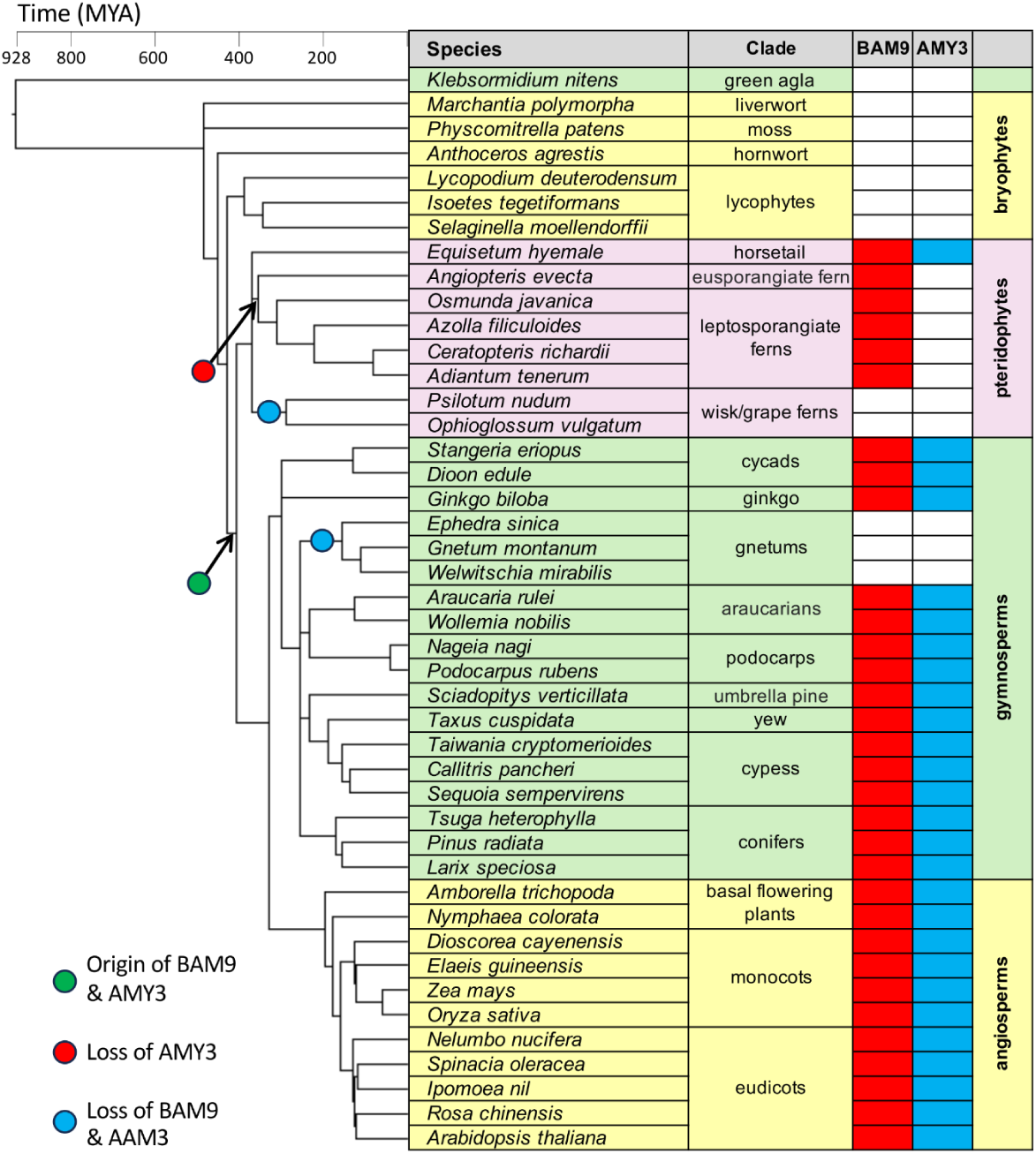
Evolution of BAM9 and AMY3 in Viridiplantae. The presence or absence of *BAM9* (red) and *AMY3* (blue) genes mapped onto a phylogenetic tree of species ranging from a charophyte alga to eudicots. The green circle indicates the putative origin of both genes. The red circle indicates a loss of the *AMY3* gene, and the two blue circles indicate the loss of both *BAM9* and *AMY3* genes. The phylogenetic tree was generated using TimeTree: timetree.org (Kumar et al. 2017) and the scale bar represents time in millions of years. Accession numbers are listed in Supplementary Table S4.

## DISCUSSION

BAM9 is a widely distributed, conserved member of the β-amylase family in plants with a degenerate and non-catalytic active site. Although the functional role of BAM9 is unclear, it has an observed effect on starch degradation (David et al., 2021). Through a yeast two-hybrid screen for BAM9 binding proteins, we identified the ɑ-amylase AMY3 as a likely interacting partner due to multiple, independent prey isolations supported by AMY3’s known chloroplast-localization and function in starch degradation. Activity assays with purified recombinant proteins showed that BAM9 enhanced the starch-hydrolyzing activity of AMY3 at a ∼1:1 molar ratio and we observed a physical interaction of the proteins in crosslinked samples. We then proposed that BAM9 forms a heterodimer with AMY3 based on solution MALS and SAXS analysis. Interestingly, we found that AMY3 also exists as a homodimer in solution, which had not previously been observed. We hypothesize that in the absence of BAM9, AMY3 forms a homodimer that has low activity. The presence of BAM9 disrupts the AMY3 homodimer, and the resulting AMY3-BAM9 heterodimer has increased activity. Whether BAM9 is necessary for AMY3 activity *in vivo* or just enhances its activity is still unknown. Recombinant AMY3 is somewhat unstable, and partial degradation of the enzyme leads to a decline in self-inhibition, which would be observed as a decrease in the enhancement of AMY3 activity by BAM9.

AMY3 has been reported to facilitate starch degradation in response to abiotic stress (Thalmann et al., 2016). However, the available transcriptomic data indicate that there is no strong influence of abiotic stress on the expression of AMY3 (Winter et al., 2007). BAM9, in contrast, does appear to be regulated at the transcriptional level by abiotic stress (Baena-González et al. 2007; Viana et al., 2021). Our results show that AMY3 activity is at least influenced by the presence of BAM9, which may explain why AMY3 is not strongly regulated by stress, as BAM9 could be expressed and enhance AMY3 activity in response to abiotic stress. Plants lacking BAM9 were found to accumulate a small excess of starch, but in combination with a BAM3 mutation and especially mutations in both BAM3 and BAM1, more starch accumulated (David et al., 2021). In the absence of both active BAMs, plants would likely rely more on AMY3 to degrade starch. If our model of the function of BAM9 activating AMY3 is correct, then plants lacking BAM9 and both BAM1 and BAM3 would be expected to accumulate more starch than plants lacking just the two active BAMs.

Our structural analysis and modeling have revealed a possible mechanism for the influence of BAM9 on AMY3. The apparent competition of BAM9 for AMY3 suggests that BAM9 disrupts the AMY3 dimer and replaces one of the AMY3 monomers in the dimer. Our structural models of the AMY3 homodimer suggest the CBM of AMY3 is proximal to the amylase domain, possibly blocking substrate access to one or both domains. The presence of BAM9 displaces one of the AMY3 proteins, increasing activity by freeing the CBM and/or the active site of AMY3 to bind to the substrate. Whether the BAM9-AMY3 complex is the active form of the enzyme or if the displaced AMY3 is now free to cleave polysaccharide chains is not clear from the present data. In either scenario, BAM9 is clearly inactive toward starch substrates and regulates the activity of an active amylase consistent with the class of proteins called pseudoenzymes. To our knowledge, this is the first definitive description of a pseudoamylase.

The pseudoamylase function of BAM9 appears to co-occur with the alpha-alpha hairpin structure of AMY3 based on the co-occurrence of these proteins (and AF3 modeling of homolog pairs) across many sequenced vascular plants. Suggestions that this AAH structure interacts with BAM9 came from the yeast two-hybrid data in which 3 unique, independent interacting prey plasmids included this region, AlphaFold3 modeling, which is based in part on coevolution of sites, and the finding that AMY3 lacking the N-terminal region (tAMY3 or AMY3_cat_) were unresponsive to BAM9. The CBM of AMY3 are known to bind carbohydrates *in vitro* and *in vivo* and are important for localization to starch, making this domain less likely to be the interaction site with BAM9 (Glaring et al., 2011).

Pseudoenzyme regulation of amylase activity is increasingly becoming apparent. LSF1, a pseudo-phosphatase, has been shown to interact with amylases BAM1, BAM3, and AMY3 (Schreier *et al*., 2019). Recent work shows that LSF1 can increase the apparent BAM1 activity by several-fold (Schreier *et al*., 2019; Liu *et al*., 2023). The authors of this latter study proposed that LSF1 feeds the starch substrate into the BAM1 active site, analogous to the CBMs found in bacterial BAMs (Liu *et al*., 2023). We find this role implausible for BAM9 given that BAM9 shows no/low affinity for starch *in vitro* (David *et al*., 2021). However, interaction between AMY3 and BAM9 may release the AMY3 CBMs from the amylase domain allowing for more efficient substrate binding. The behavior of BAM9 is consistent with the regulatory actions taken by pseudoenzymes from other families (Eyers and Murphy, 2016; Jeffery, 2019). Thus we classify BAM9 as a pseudoamylase regulating starch degradation by AMY3 in response to abiotic stress.

Our work is the first description of a pseudoamylase and provides a framework for assessing the other inactive BAMs found in Arabidopsis and other plant species (ex. Arabidopsis BAM4). BAM4, like BAM9, is found in the chloroplast and lacks the amino acids required for starch hydrolysis but BAM4 knockouts show a negative effect on starch degradation (Fulton *et al*., 2008; Comparot-Moss *et al*., 2010). These findings are consistent with the behaviors of pseudoenzymes and suggest pseudoamylases regulate many aspects of starch degradation, although the specific mechanisms are unclear.

## Acknowledgments

We thank Kevin Fedkenheuer, John Herlihy, Isabella Law, Valeria Moscote-Rodriguez, Nurlybeck Mursaliyev, Samantha Platt, Lauren Saunders, Karen Siddoway, Elizabeth Steidle, and Kitrina Wargo, who did early work on BAM9 and AMY3, which ultimately led to the current research direction. This work was supported by NSF IOS RUI-1146776, MCB RUI-1616467, MCB RUI-1932755, MCB RUI-2322867, and CHE REU-2150091. This work was conducted in part at the Advanced Light Source (ALS), a national user facility operated by Lawrence Berkeley National Laboratory on behalf of the Department of Energy, Office of Basic Energy Sciences, through the Integrated Diffraction Analysis Technologies (IDAT) program, supported by DOE Office of Biological and Environmental Research. Additional support comes from the National Institute of Health project ALS-ENABLE (P30 GM124169) and a High-End Instrumentation Grant S10OD018483.

## Notes

### Competing Interest Statement

The authors have declared no competing interest.

