## Supplemental File for "The pseudoenzyme β-amylase9 from Arabidopsis binds to and enhances the activity of α-amylase3: A possible mechanism to promote stress-induced starch degradation"

**Supplementary Table S1.** Summary of yeast-two hybrid prey plasmids isolated with a BAM9 bait protein that encode predicted chloroplast proteins.

**Supplementary Table S2.** BAM9 and AMY3 protein sequence identification numbers used for WebLogo construction.

**Supplementary Table S3.** SAXS data collection and fitting statistics.

**Supplementary Table S4.** BAM9 and AMY3 protein sequence identification numbers used for phylogenetic analysis.

**Supplementary Figure S1.** Effect of BAM9 on AMY3 activity toward various substrates and the effect of salts on AMY3 activity.

**Supplementary Figure S2.** Predicted aligned error plot for AlphaFold3 prediction of Arabidopsis AMY3 and BAM9.

**Supplementary Figure S3.** Top scoring AlphaFold3 models of BAM9-AMY3 complexes from representative vascular plants.

**Supplemental Table 1.** Summary of yeast-two hybrid prey plasmids isolated with a BAM9 bait protein that encode predicted chloroplast proteins.

| (Screen number) and Isolate Label | Number of Clones | Gene Number | UniProt Identifier | Protein Name |
|---|---|---|---|---|
| (1) 9B4 | 1 | AT2G06520 | Q9SKI3 | Photosystem II subunit PsbX |
| (1) 10A2 | 1 | AT1G10095 | O80601 | Protein prenylyltransferase superfamily protein |
| (1)14A3 | 1 | AT1G29930 | P04778 | Chlorophyll a-b binding protein 1 |
| (1) 17A4 | 1 | AT4G19170 | O49675 | Carotenoid cleavage dioxygenase 4 |
| (2) 1A1, 7A2, 6A3, 8A2 | 4* | AT1G69830 | Q94A41 | Alpha-amylase 3 |
| (2) 1B2, | 2 | AT1G13270 | Q9FV52 | Methionine aminopeptidase 1B |
| (2) 20B1 | 1 | AT3G63160 | Q9M1X3 | Outer envelope protein 6; OEP 7.2 |
| (2) 5A2 | 1 | AT1G11870 | Q8RWT8 | Seryl-tRNA synthetase |

* Two isolates had the same sequence but all were isolated independently.

**Supplemental Table 2.** BAM9 and AMY3 protein sequence identification numbers used for WebLogo construction. Twenty-one different orders were included.

| Order | Species | BAM9 ID#(s) | |
|---|---|---|---|
| Proteales | *Nelumbo nucifera* | XP_010241169.1 | XP_010256483.1 |
| Caryophyllales | *Spinacia oleracea* | XP_021862263.1 | XP_021844627.1 |
| Nymphaeales | *Nymphaea colorata* | XP_031498066.1 | XP_031504981.1 |
| Zingiberales | *Musa acuminata* | XP_064976691.1 | XP_009412382.1 |
| Arecales | *Elaeis guineensis* | XP_010938702.1 | XP_010925057.1 |
| Poales | *Zea mays* | NP_001170007.1<br>XP_008660155.1 | XP_008675104.1 |
| Amborallales | *Amborella trichopoda* | XP_006855410.1 | XP_011622500.1 |
| Dioscoreales | *Dioscorea cayenensis* | XP_039136687.1 | XP_039117379.1 |
| Magnoliales | *Magnolia sinica* | XP_058077990.1 | XP_058105304.1 |
| Brassicales | *Arabidopsis thaliana* | NP_197368.1 | NP_564977.1 |
| Malvales | *Gossypium hirsutum* | XP_016697781.2 | XP_016705150.2 |
| Solanales | *Ipomoea triloba* | XP_031112264.1 | XP_031124848.1 |
| Malpighiales | *Ricinus communis* | XP_002516865.1 | XP_002520134. |
| Rosales | *Rosa chinensis* | XP_024197540.1 | XP_024195098.1 |
| Cucurbitales | *Cucumis sativus* | XP_004144975.1 | XP_004135194.1 |
| Vitales | *Vitis vinifera* | XP_002276777.1 | XP_002270049.1 |
| Apiales | *Daucus carota* | XP_017219710.1 | XP_017222730.1 |
| Fabales | *Medicago truncatulata* | XP_003594004.1 | XP_013445797.1 |
| Ericales | *Camellia sinensis* | XP_028061385.1 | XP_028099207.1 |
| Lamiales | *Sesamum indicum* | XP_011090854.1 | XP_011086391.1 |
| Asterales | *Lactuca sativa* | XP_023758663.1 | XP_052623018.1 |

### Supplemental Table 3 – SAXS data collection and fitting statistics

| | BAM9 | AMY3 | BAM9-AMY3 |
|---|---|---|---|
| **Wavelength (Å)** | 1.127 | 1.127 | 1.127 |
| **q range (Å-1)** | .0099 - .4734 | .0099 - .4734 | .0099 - .4734 |
| **Concentration (mg mL-1)** | 4 | 4 | 4 |
| **Temperature (K)** | 283 K | 283 K | 283 K |
| **I(0) (A.U.) [from p (r )]** | 40.46 +/- 0.13 | 87.03 +/- 1.4 | 89.99 +/- 1.48 |
| **Rg (Å) [from p (r )]** | 24.57 +/-0.11 | 55.76 +/- 1.01 | 65.22 +/- 1.1 |
| **I(0) (A.U.) [from Guinier]** | 40.33 +/- 0.12 | 83.92 +/- 1.7 | 77.67 +- 1.61 |
| **Rg (Å) [from Guinier]** | 24.36 +/- 0.22 | 50.96 +/- 1.18 | 50.47 +/- 1.19 |
| **Dmax (Å)** | 87 | 215 | 255 |
| **Porod volume estimate (Å3) (Vp)** | 55000 | 249000 | 212000 |
| **Molecular mass Bayes (kDa)** | 42 | 169.6 | 146.8 |
| **Molecular mass Mr [from Porod volume] (kDa)** | 45.7 | 206.3 | 175.7 |
| **Calculated monomeric Mr from sequence (Da)** | 50300 | 93600 | 143900 |
| **SASBDB ID** | | | |
| **Simple Scattering dataset code** | XS1RR5P9 | XSVTUF83 | XS5XWKRW |
| **Chi^2 FOXS fit to model** | 1.02 | 0.8 | |
| **Primary data reduction** | RAW | RAW | RAW |
| **Data processing** | RAW | RAW | RAW |
| **Rigid body modeling** | | BilboMD | |
| **Computation of model intensities** | FOXS | FOXS | |
| **Three-dimensional graphic representations** | YASARA | YASARA | |

**Table S4.** BAM9 and AMY3 protein sequence identification numbers used for phylogenetic analysis.

| Species | Common name | BAM9 ID# | AMY3 ID# |
|---|---|---|---|
| *Equisetum hyemale* | horsetail | JVSZ_scaffold_2015189, JVSZ_scaffold_2131870 | JVSZ_scaffold_2132479 |
| *Angiopteris evecta* | eusporangiate fern | NHCM_scaffold_2010456 | - |
| *Osmunda javanica* | leptosporangiate ferns | VIBO_scaffold_2079058 | - |
| *Azolla filiculoides* | | Azfi_s0540.g076343 | - |
| *Ceratopteris richardii* | | PIVW_scaffold_2013454 | - |
| *Adiantum tenerum* | | BMJR_scaffold_2015754 | - |
| *Stangeria eriopus* | cycads | KAWQ_scaffold_2056927 | KAWQ_scaffold_2010767 |
| *Dioon edule* | | WLIC_scaffold_2121558 | WLIC_scaffold_2121748 |
| *Ginkgo biloba* | ginkgo | Gb_27777 | Gb_02250 |
| *Araucaria rulei* | araucarians | XTZO_scaffold_2066135 | XTZO_scaffold_2002814 |
| *Wollemia nobilis* | | RSCE_scaffold_2059177 | RSCE_scaffold_2060518 |
| *Nageia nagi* | podocarps | UUJS_scaffold_2116664 | UUJS_scaffold_2117889 |
| *Podocarpus rubens* | | XLGK_scaffold_2011763 | XLGK_scaffold_2009754 |
| *Sciadopitys verticillata* | umbrella pine | YFZK_scaffold_2009143 | YFZK_scaffold_2009076 |
| *Taxus cuspidata* | yew | ZYAX_scaffold_2075462 | ZYAX_scaffold_2011158 |
| *Taiwania cryptomerioides* | cypress | QSNJ_scaffold_2068652 | QSNJ_scaffold_2069520 |
| *Callitris pancheri* | | JDQB_scaffold_2002769 | JDQB_scaffold_2010916 |
| *Sequoia sempervirens* | | HBGV_scaffold_2049744 | HBGV_scaffold_2002007 |
| *Tsuga heterophylla* | conifers | GAMH_scaffold_2057735 | GAMH_scaffold_2009337 |
| *Pinus radiata* | | DZQM_scaffold_2055847 | DZQM_scaffold_2056253 |
| *Larix speciosa* | | WVWN_scaffold_2055349 | WVWN_scaffold_2055893 |
| *Amborella trichopoda* | basal flowering plants | XP_006855410.1 | XP_011622500.1 |
| *Nymphaea colorata* | | XP_031498066.1 | XP_031504981.1 |
| *Dioscorea cayenensis* | monocots | XP_039136687.1 | XP_039117379.1 |
| *Elaeis guineensis* | | XP_010938702.1 | XP_010925057.1 |
| *Zea mays* | | NP_001170007.1 XP_008660155.1 | XP_008675104.1 |
| *Oryza sativa* | | XP_015646194.1 XP_015631450.1 | XP_015649620.1 |
| *Nelumbo nucifera* | eudicots | XP_010241169.1 | XP_010256483.1 |
| *Spinacia oleracea* | | XP_021862263.1 | XP_021844627.1 |
| *Ipomoea triloba* | | XP_019172872.1 | XP_031124848.1 |
| *Rosa chinensis* | | XP_024197540.1 | XP_024195098.1 |
| *Arabidopsis thaliana* | | NP_197368.1 | NP_564977.1 |

Supplemental Figure 1

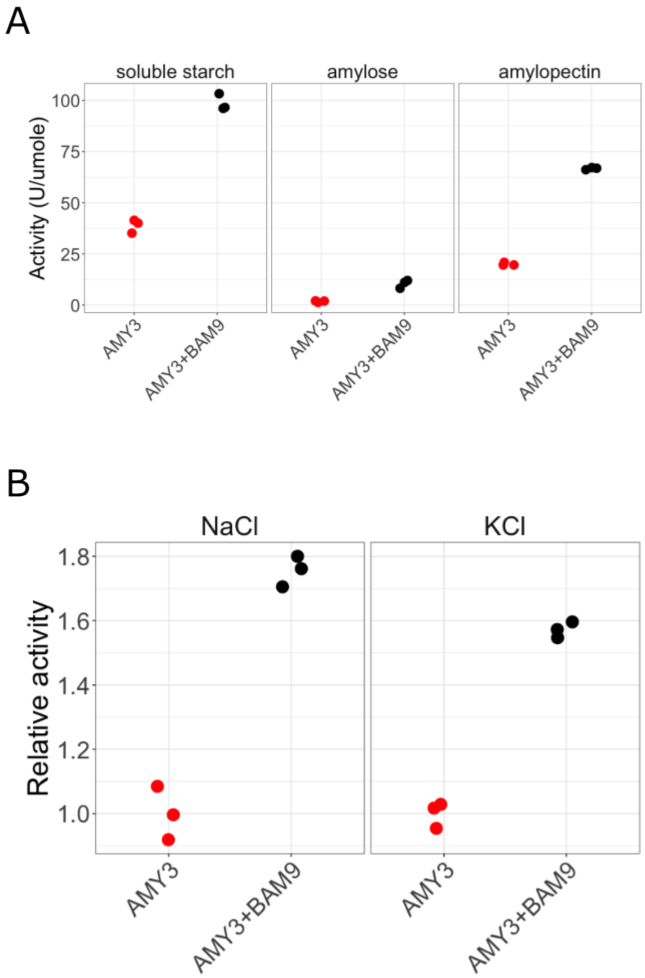

(A) Activity of AMY3 and AMY3+BAM9 on soluble starch, amylose, and amylopectin. Data shown are from three activity assays. (B) Effect of 100 mM NaCl or 100 mM KCl on activity of AMY3 and AMY3+BAM9. Data were normalized to the respective level of the AMY3 reaction. Data from three activity assays are shown.

Supplemental Figure 2

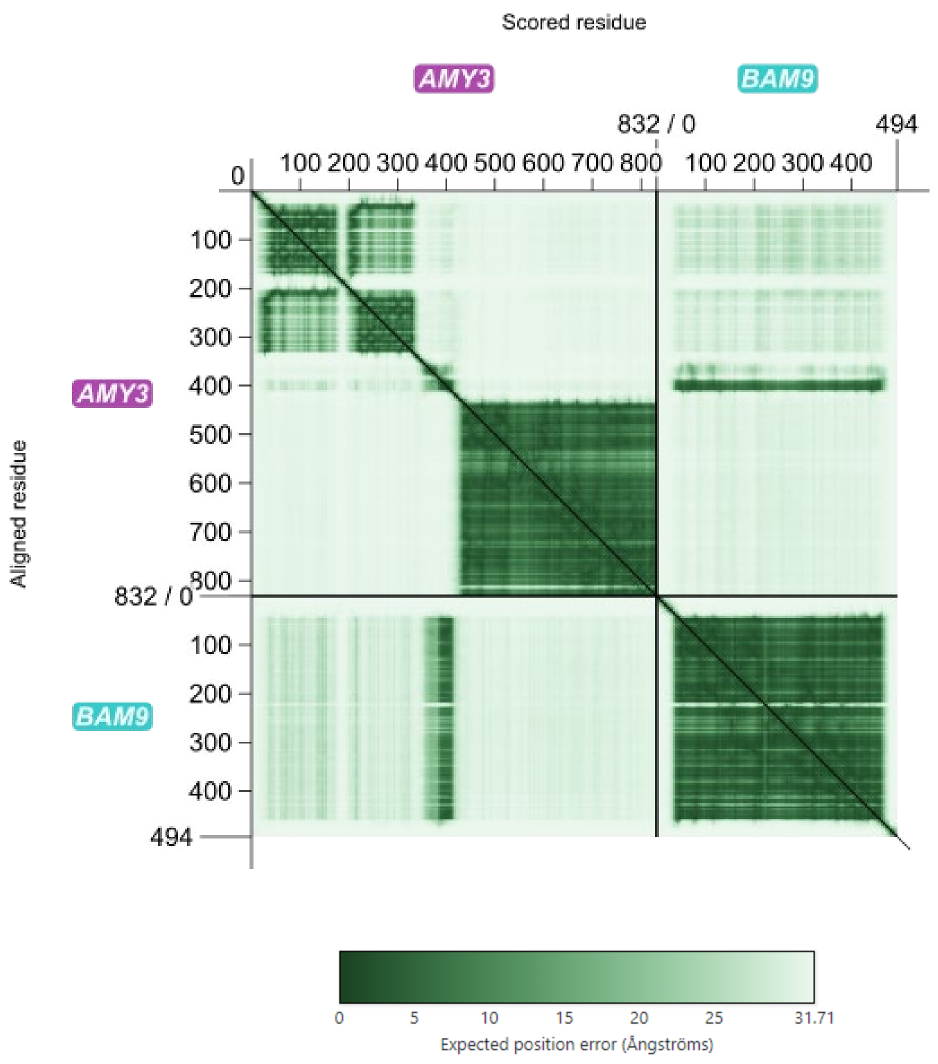

Predicted aligned error plot for AlphaFold3 prediction of AMY3 and BAM9.

Supplemental Figure 3

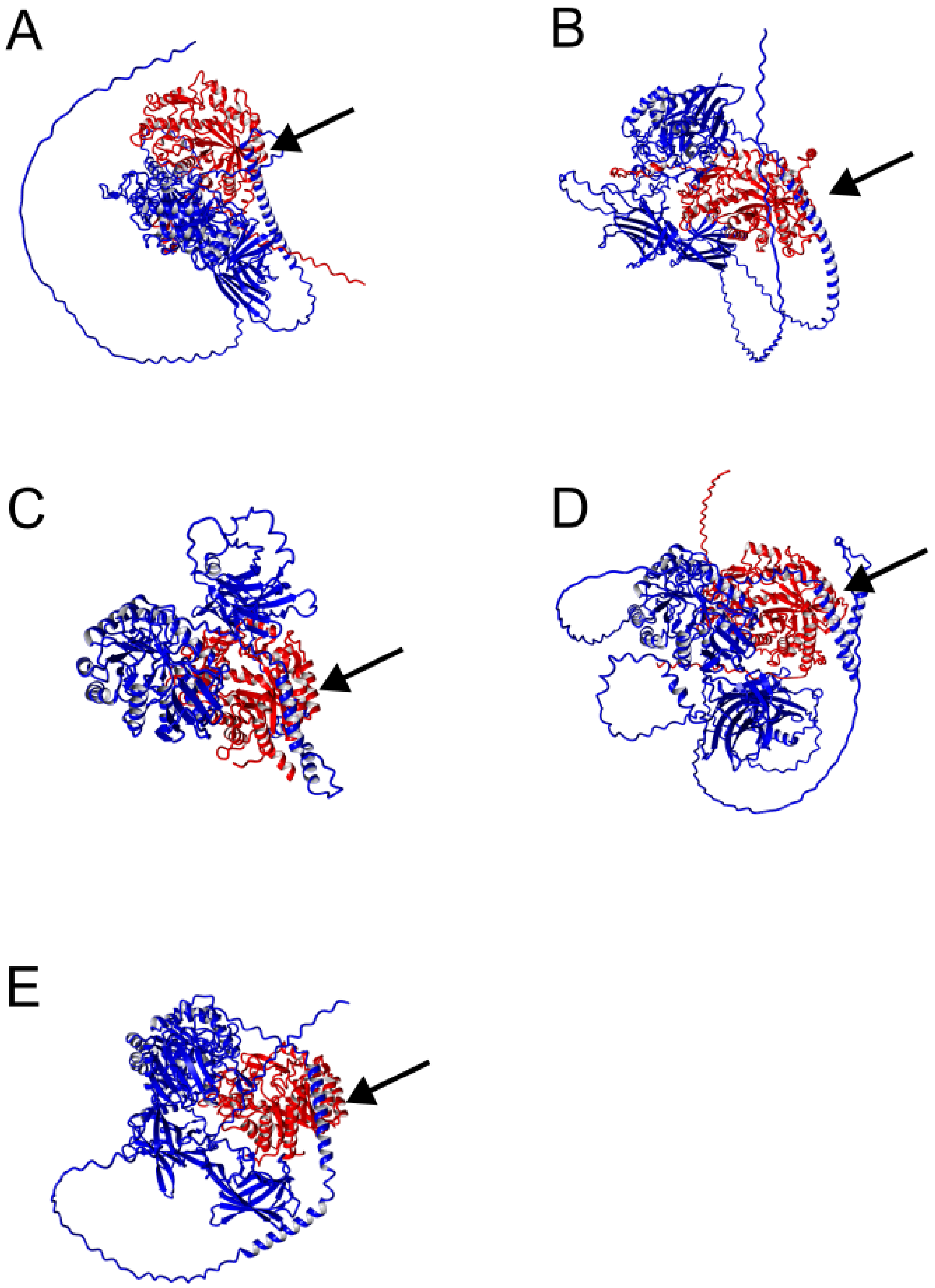

Top scoring AlphaFold3 models of BAM9-AMY3 complexes from (A) and (B) *Zea mays* (C) Equisetum (D) Gingko, and (E) *Pinus radiata*. *Zea mays* has two potential BAM9 homologs, both of which appear to be able to bind to AMY3 from that plant. The arrow in each panel highlights the alpha-alpha hairpin of AMY3 and BAM9 interaction site in each model.
